# Antibacterial Activity of *Andrographis paniculata* and *Piper betle* and their Interactive Effects with Amoxicillin Against Selected Respiratory Pathogens

**DOI:** 10.1101/2023.04.05.535803

**Authors:** Cristian B. Mejos, Joshua G. Poblete, Paula Jean C. Sarino, Ma. Fatima I. Cruzada

## Abstract

This study was undertaken to determine the antibacterial activity and interactive effects of the methanol, ethanol and aqueous extract of *Andrographis paniculata* and *Piper betle* leaves with amoxicillin against selected clinical isolates of respiratory pathogens: *Escherichia coli* USTCMS 1030, *Pseudomonas aeruginosa USTCMS* 10013, and *Staphylococcus aureus* USTCMS 1097. Antibacterial activity of the plant extracts using disk diffusion showed that the methanol extract of *P. betle* exhibited inhibitory activity against all the test organisms, whereas the methanol and ethanol extracts of *A. paniculata* exhibited antibacterial activity to *S. aureus* USTCMS 1097 only. The antimicrobial properties of each plant extract were further evaluated using broth microdilution. Results showed that the ethanol extract of *P. betle* had the most potent antibacterial activity against all test bacteria with minimum inhibitory concentrations of 6.5 mg/mL, 3.25 mg/mL, and 0.2 mg/mL for *E. coli* USTCMS 1030, *P. aeruginosa* USTCMS 10013 and *S. aureus* USTCMS 1097, respectively. However, resazurin showed an inhibitory activity against *S. aureus* USTCMS 1097 in usual concentrations used in the assay, which is a novel finding since it is typically used as an indicator. Based on disk diffusion, the methanol and aqueous extracts of *P. betle* showed promising synergistic effect with the antibiotic amoxicillin. This was confirmed by checkerboard assay wherein the aqueous extract of *P. betle* showed an additive effect to amoxicillin against *E. coli* USTCMS 1030 (FICI = 0.66), while the methanol extract of *P. betle* exhibited true synergism with amoxicillin against *P. aeruginosa* USTCMS 10013 (FICI = 0.33). This synergism between the ethanol extract of *P. betle* and amoxicillin was significant since the activity of amoxicillin increased by 128-fold. This combination has potential in treating diseases associated with amoxicillin-resistant *P. aeruginosa*.

## INTRODUCTION

Lower respiratory tract infections (LRTIs) are infections affecting the pneumonic system of humans consisting of tuberculosis, pneumonia, and bronchitis (Fletcher, 2019). It is a considerable culprit of respiratory illnesses and deaths persistent in intensive care units (ICU) (Cardoso, 2007). This accounts for nearly 4 million deaths annually, affecting children under 5 years of age up to those people 70 years and older (Trouger, 2017; Forum of International Respiratory Societies, 2017).

Bacteria are known to cause some of the diseases associated with LRTIs, wherein some of the frequent causal agents isolated from the patients were *Escherichia coli*, *Staphylococcus aureus*, and *Pseudomonas aeruginosa* (Noviello & Huang, 2019). In addition, bacterial co-infections of strains mentioned were observed in the respiratory tract of critically-ill Coronavirus Disease 2019 (COVID-19) patients in the ICU (Prasetyoputri, 2021).

LRTIs are commonly treated with antibiotics (Kraus *et al*., 2017), such as amoxicillin. When used correctly, antibiotics can inhibit the growth of specific pathogens, or kill them (Anderson, 2021). However, the widespread abuse and misuse 2 of antibiotics has caused the development and spread of antibiotic resistance in bacteria, making many bacterial infections difficult to treat (Dugassa & Shukuri, 2017). No new antibiotics were produced for more than 35 years already, and so there are limited antimicrobial arsenals that can be utilized for treating diseases (Perez, 2021). Hence, a new approach for treating these diseases must be explored.

This issue was intensified by the recent pandemic since COVID-19 patients most often receive antibiotic medications. This further increases the concentration of antibiotics in the environment which can promote the rise of resistant pathogens (Knight *et al*., 2021). The acquired drug resistance of bacteria to existing antibiotics obliges scientists to look for new antimicrobial agents that can address this issue (Fair & Tor, 2014). This led to the introduction of other integrants such as plants that had been in consideration evidential from recent studies.

Plants offer a cheap source of chemical entities and plant-derived compounds, which exhibit antimicrobial properties. In fact, a quarter of all the antibiotics in the market today were derived from plant-based products (Abreu *et al*., 2012). Herbal plants like the *Andrographis paniculata* and *Piper betle* are traditionally used for the treatment and medication of various diseases, particularly in an Asian country like the Philippines. These medicinal plants have been used way before the advancement of modern medicine and are still being used as an alternative form of health care (Valle, 2015; Okhuarobo *et al*., 2014). *A. paniculata* has antifungal (Sule *et al*., 2012), antiprotozoal (Okhuarobo *et al*., 2014), and antibacterial properties (Arifullah *et al*., 2013), while *P. betle* has antibacterial, antioxidant, anticarcinogen (Prakash *et al*., 2010), antifungal (Ali *et al*., 2010), and antiparasitic properties (Pecková *et al*., 2018).

Although there have been numerous scientific studies about the antimicrobial property of *A. paniculata* and *P. betle* against different microorganisms, there has never been any research about their effects against *E. coli*, *S. aureus*, and *P. aeruginosa* that are associated with LRTIs. Also, data on interactive effects of these 3 herbal plants combined with amoxicillin are limited. Hence, this study sought to assess the antimicrobial activity of these plant extracts, mainly, *A. paniculata* and *P. betle* as well as studying their interactive effects when combined with amoxicillin against selected respiratory pathogens.

## RESULTS AND DISCUSSION

### Extraction Yield for Each Plant Sample

Different solvents were used for plant extraction and these included water, ethanol, and methanol. The utilization of different solvents resulted in the extraction of different potential antimicrobial compounds of *Andrographis paniculata* and *Piper betle*. The methanol and ethanol extracts of *A. paniculata* appeared to have higher extraction yield compared to the *P. betle* that was extracted using the same solvents (Table 1). On the other hand, for the aqueous extract, a much higher yield was obtained from the *P. betle* compared to that of the *A. paniculata*. The yields obtained from *A. paniculata* in this study were comparatively lower than those obtained by Kumoro *et al*. (2009). For *P. betle*, the yield results were similar to those obtained by Azahar *et al*. (2023) wherein the methanol extract had higher yield than the ethanol extract. The extraction yield of *P. betle* in the present study is also similar to the study of Foo’s team (2015), where the aqueous extract had a higher yield compared to the ethanol extract. Overall, in *P. betle*’s case, the yield of ethanol extract was always lesser than the methanol and aqueous extracts. The difference in the extraction yield of this study compared to other studies may possibly be due to the difference in the extraction and purification methods since some studies employed the use of a Soxhlet extraction system.

**Table 1.** Extraction yield and concentration of A. paniculata and P. betle.

| PLANT SPECIES | EXTRACTION YIELD (%) | EXTRACT CONCENTRATION (mg/mL) |
| --- | --- | --- |
| *Andrographis paniculata* |  |  |
| Methanol Extract | 18.55 | 463.75 |
| Ethanol Extract | 8.57 | 244.95 |
| Aqueous Extract | 3.378 | 225.20 |
| *Piper betle* |  |  |
| Methanol Extract | 8.45 | 281.70 |
| Ethanol Extract | 1.04 | 52.00 |
| Aqueous Extract | 10.33 | 688.6 |

### Antimicrobial Activity of Plant Extracts using Disk Diffusion Assay

The antimicrobial activity of methanol, ethanol and aqueous extracts of *A. paniculata* and *P. betle* against *Escherichia coli* USTCMS 1030, *Pseudomonas aeruginosa* USTCMS 10013, and *Staphylococcus aureus* USTCMS 1097 were initially assessed using the disk diffusion assay (Table 2). Only the methanol extract of *P. betle* was observed to have shown an inhibition zone against *E. coli* USTCMS 1030 (Figure 1). All other plant extracts did not have any visible inhibition zone against *E. coli* USTCMS 1030. As for the amoxicillin, it had a higher zone of inhibition with an average of 19 mm compared to the methanol extract of *P. betle*. These results disagree with the previous data obtained in a study by Paderes and Bose (2020) that the ethanol extract of *A. paniculata* had a moderate inhibitory activity against *E. coli* USTCMS 1030.

**Table 2.** Antimicrobial activity of plant extracts against selected pathogens using the disk diffusion assay.

| TEST STRAIN | ZONE OF INHIBITION (mm) / EXTRACT |  |  |  |  |  | *Antibiotic |
| --- | --- | --- | --- | --- | --- | --- | --- |
|  | Andrographis paniculata |  |  | Piper betle |  |  |  |
|  | Methanol | Ethanol | Aqueous | Methanol | Ethanol | Aqueous |  |
| Escherichia coli<br>USTCMS 1030 | 0 | 0 | 0 | 12.5 ± 0.5<sup>b</sup> | 0 | 0 | 19 ± 0.2<sup>a</sup> |
| Pseudomonas aeruginosa<br>USTCMS 10013 | 0 | 0 | 0 | 7.66 ± 0.6<sup>a</sup> | 0 | 0 | 0 |
| Staphylococcus aureus<br>USTCMS 1097 | 7.75 ± 0.3<sup>b</sup> | 7.5 ± 0.3<sup>b</sup> | 0 | 8.83 ± 0.2<sup>b</sup> | 0 | 0 | 28.5 ± 0.2<sup>a</sup> |

Values are means of triplicate determination ( $n = 3$ ) ± Standard Error of the Mean (SEM); \*Antibiotic values are also in means but are computed from nine data measurements ( $n = 9$ ). Lowercase letters in the same row indicate significant differences ( $P < 0.05$ ).

On the other hand, the result for aqueous extract was similar with Zaidan *et al*. (2005) where the extract did not exhibit any inhibitory activity against *E. coli* ATCC 25922 at all concentrations. In another previous study, clinical isolates of *E. coli* were also subjected to ethanol and methanol extracts of *P. betle*, and they showed antimicrobial activity through disk diffusion assay (Valle *et al*., 2016). However, in the present study, only the methanol extract of *P. betle* produced an antimicrobial activity with an inhibition zone of 12.5 ± 0.5 mm.

The results of the disk diffusion assay revealed that only one out of the six extracts exhibited antimicrobial activity against *P. aeruginosa* USTCMS 10013 (Figure 2). Methanol extract of *P. betle* displayed an average inhibition zone of 7.66 ± 0.6 mm, while amoxicillin was completely resisted by *P. aeruginosa* USTCMS 10013. The results for the ethanol and methanol extracts of *P. betle* differed from the study of Zaidan *et al*. (2005) that recorded significant antimicrobial activity against *P. aeruginosa*. As for the extracts of *A. paniculata*, their results also disagree with the findings of the present study since they detected an antimicrobial activity of its water extract against *P. aeruginosa*.

**Figure 2.**
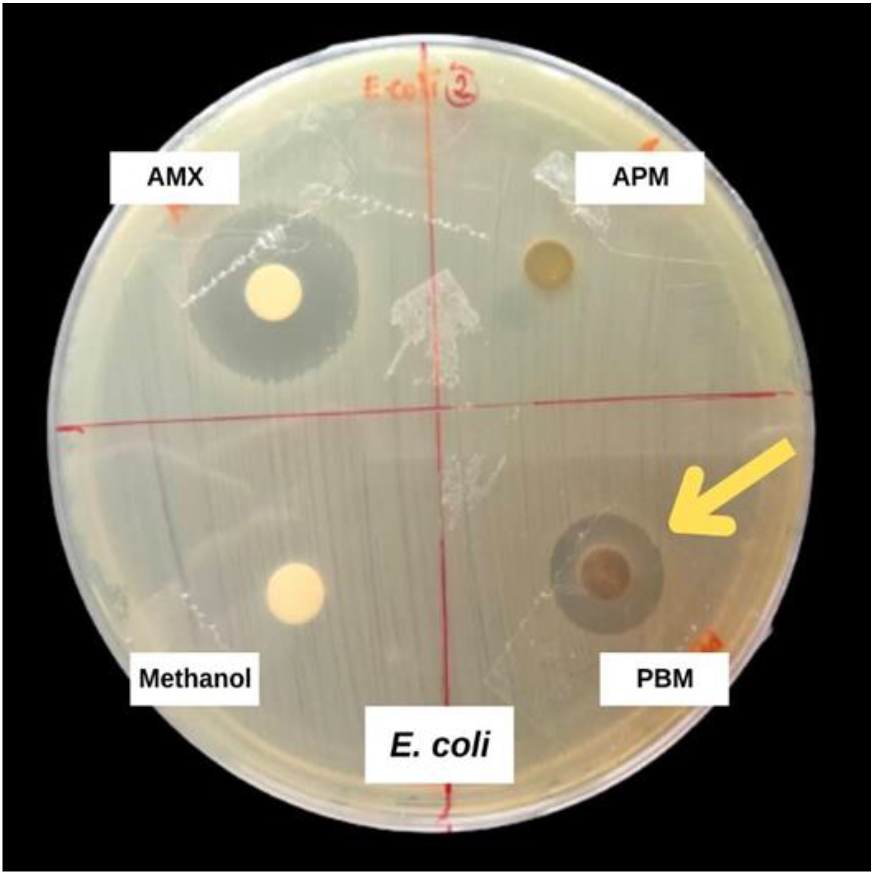
Antibacterial activity of the methanol extract of P. betle and A paniculata against E. coli USTCMS 1030. *AMX, amoxicillin; APM, methanol extract of A. paniculata; PBM, methanol extract of P. betle.

**Figure 3.**
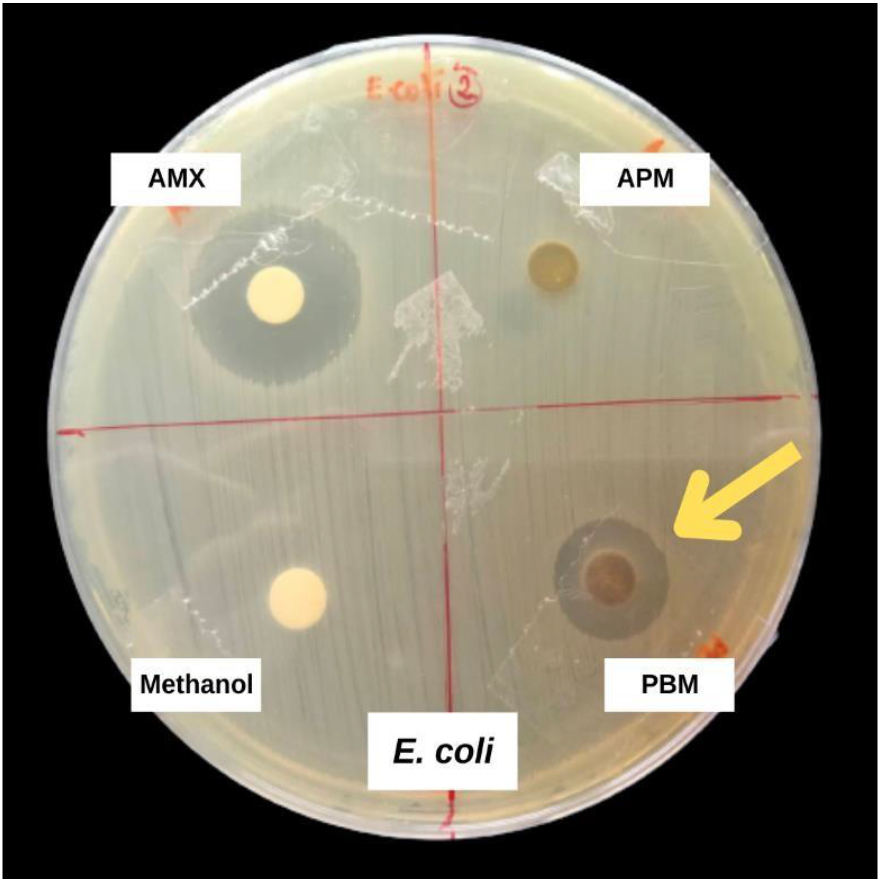
Antibacterial activity of the methanol extract of P. betle and A. paniculata against P. aeruginosa. *AMX, Amoxicillin; APM, Methanol extract of A. paniculata; PBM, Methanol extract of P. betle.

The results also revealed that both *A. paniculata* and *P. betle* had antimicrobial activities against *S. aureus* USTCMS 1097. The methanol extract of *P. betle* exhibited the highest inhibition zone (8.83 ± 0.2 mm) followed by the *A. paniculata* ethanol extract (7.5 ± 0.3 mm) and then by *A. paniculata* methanol extract (7.75 ± 0.3 mm) (Table 2). No activity was observed in *A. paniculata* aqueous extract and *P. betle* ethanol and aqueous extracts. Amoxicillin displayed the largest inhibition zone at 28.5 mm (Figure 4).

**Figure 4.**
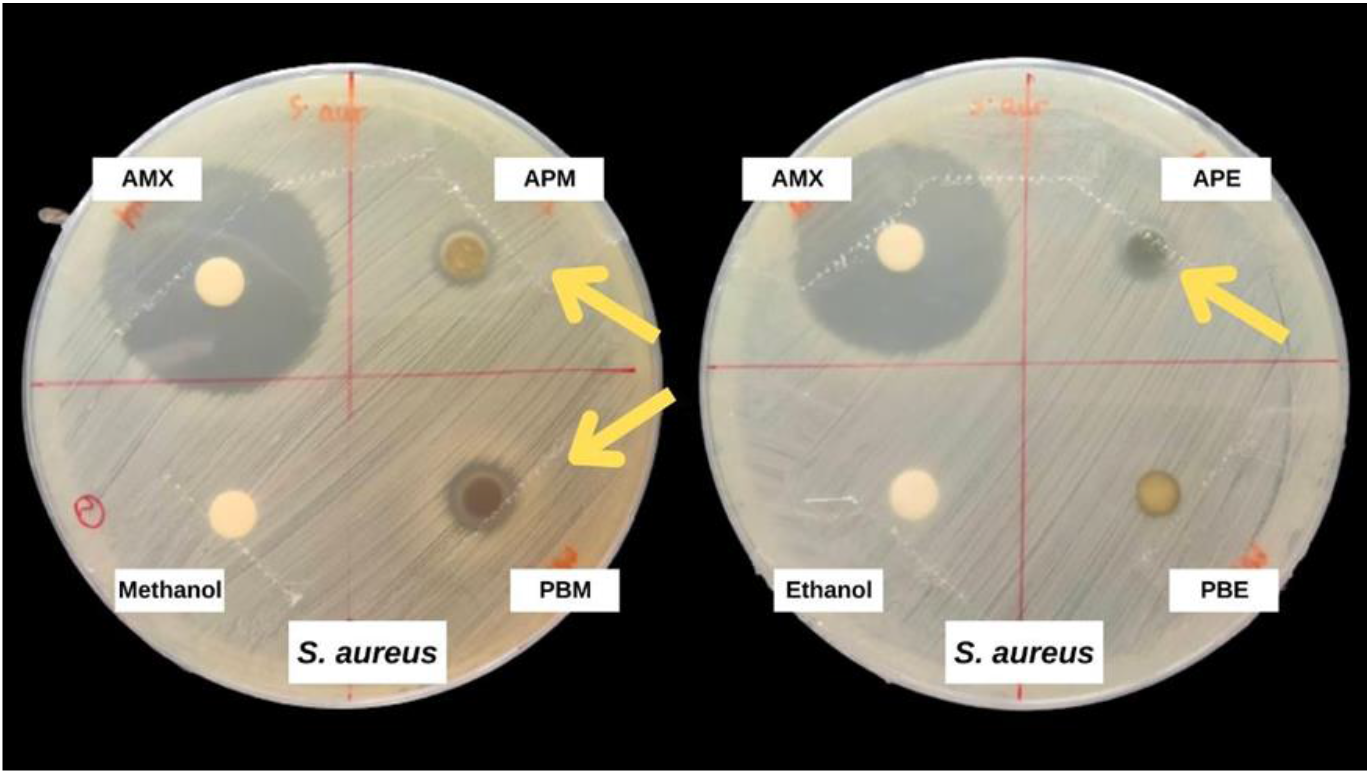
Antibacterial activity of the methanol extract of P. betle and the ethanol and methanol extracts of A. paniculata against S. aureus USTCMS 1097. *AMX, amoxicillin; APM, Methanol extract of A. paniculata; PBM, Methanol extract of P. betle; APE, Ethanol extract of A. paniculata; PBE, Ethanol extract of P. betle.

The presence of antimicrobial activity in the ethanol extract of *A. paniculata* matches with the study of Sahalan *et al*. (2007) who found moderate antibacterial activity against the *S. aureus*. *A. paniculata* aqueous extract showed weak antimicrobial activity which is similar to the results of Zaidan *et al*. (2005).

As for the methanol extract of *P. betle*, the present study is congruent with the findings of Valle *et al*. (2016) who found that many variants of Methicillin-resistant *Staphylococcus aureus* (MRSA) were inhibited by *P. betle*’s methanol extract. However, Valle’s group also detected an antimicrobial activity for the *P. betle*’s ethanol extract. Whereas in the present study, no inhibition zone was found for the ethanol extract of *P. betle* against the *S. aureus* USTCMS 1097.

The results of disk diffusion assay indicated that the methanol extract of *P. betle* has better prospects as an antibacterial agent against all the tested pathogenic strains. Moreover, methanol and ethanol extracts of *A. paniculata* were promising against *S. aureus* USTCMS 1097 compared to other extracts.

In prior years, *P. betle* and *A. paniculata* were extracted and tested for their antimicrobial activity against many pathogens such as *E. coli*, *S. aureus*, and *P. aeruginosa* using the disk diffusion method. The discrepancy in the results obtained in previous literature from the findings in the present study may possibly be due to the difference in the preparation or resuspension of the dried plant extract. Other studies made use of 0.2 % DMSO (Valle *et al*., 2016) and 10% DMSO (Sahalan *et al*., 2007) whereas the present study resuspended the pure extract in their respective solvents. It is also possible that the variation in the volume of extracts per disk affects the overall measurement of the inhibition zones since there are studies that made use of 50, 100, and 150 µL per disk (Rajalakshmi & Cathrine, 2016), while this study used 20 µL.

### Antimicrobial Activity and Minimum Inhibitory Concentration of Plant Extracts using Broth Microdilution Assay

Using the broth microdilution assay, all the extracts exhibited antimicrobial activity against the selected bacterial strains. This is in contrast to the results in the disk diffusion wherein only a few plant extracts showed antimicrobial activity. The disparity of the results of the disk diffusion assay and the broth microdilution may be due to the difference in the extract used in each assay. Also, the diffusion of bioactive compounds is relatively more difficult in the agar matrix compared to the broth. Another factor may also be the difference in the number of bacterial cells used in each assay.

The broth microdilution assay also revealed the minimum inhibitory concentration (MIC) of each plant extract (Table 3). Among all the extracts used in this study, ethanol extract of *P. betle* showed the lowest MIC values against *E. coli* USTCMS 1030 (6.5 mg/mL) (Figure 5), *P. aeruginosa* USTCMS 10013 (3.25 mg/mL) (Figure 6), and *S. aureus* USTCMS 1097 (0.2 mg/mL) (Figure 7).

**Figure 5.**
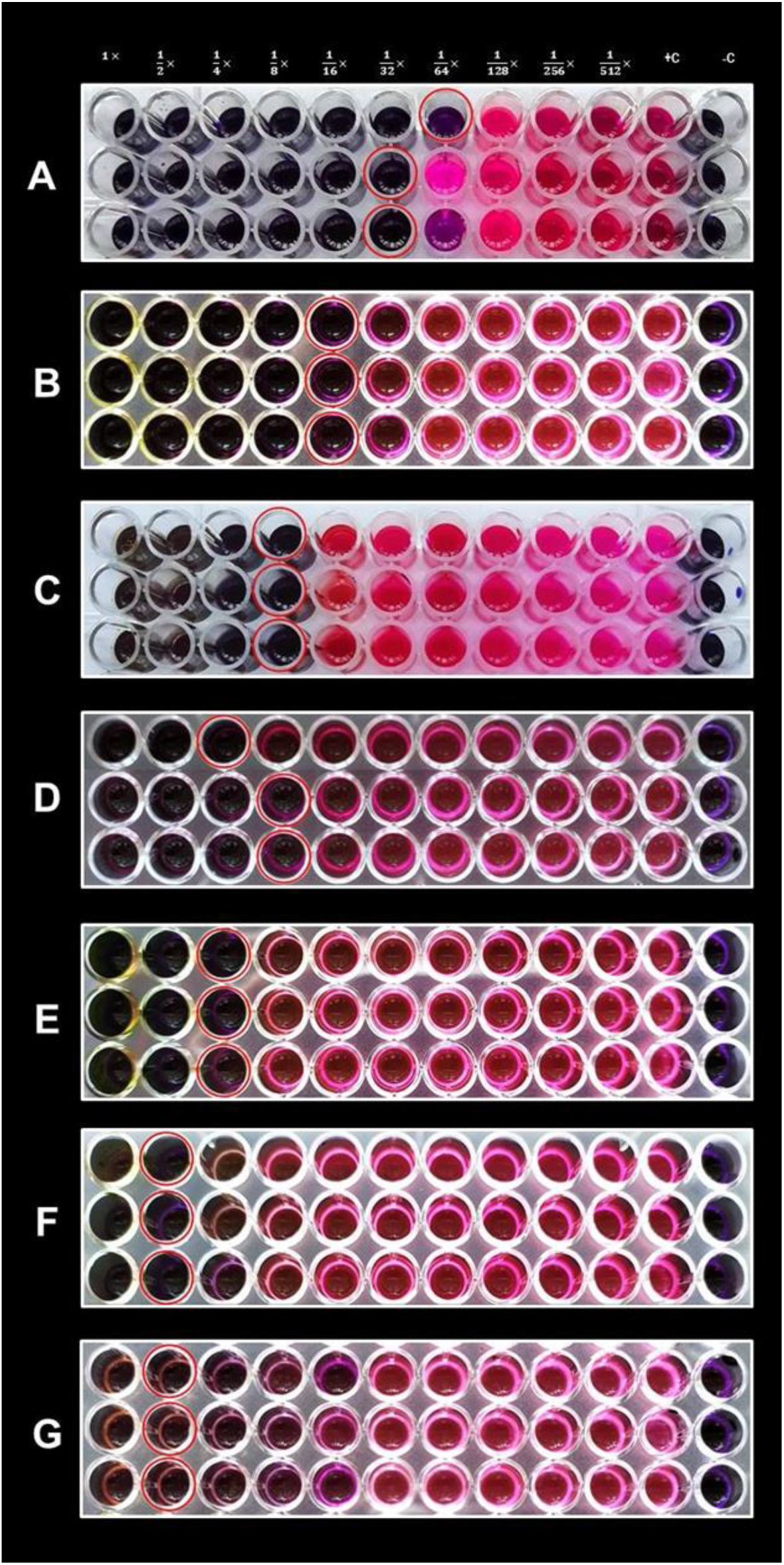
Minimum inhibitory concentrations of the plant extracts and amoxicillin against E. coli USTCMS 1030 (lowest to highest). *A, Amoxicillin; B, P. betle ethanol extract; C, P. betle methanol extract; D, A. paniculata aqueous extract; E, A. paniculata ethanol extract; F, A. paniculata methanolextract; G, P. betle aqueous extract; +c, positive control; -c, negative control

**Figure 6.**
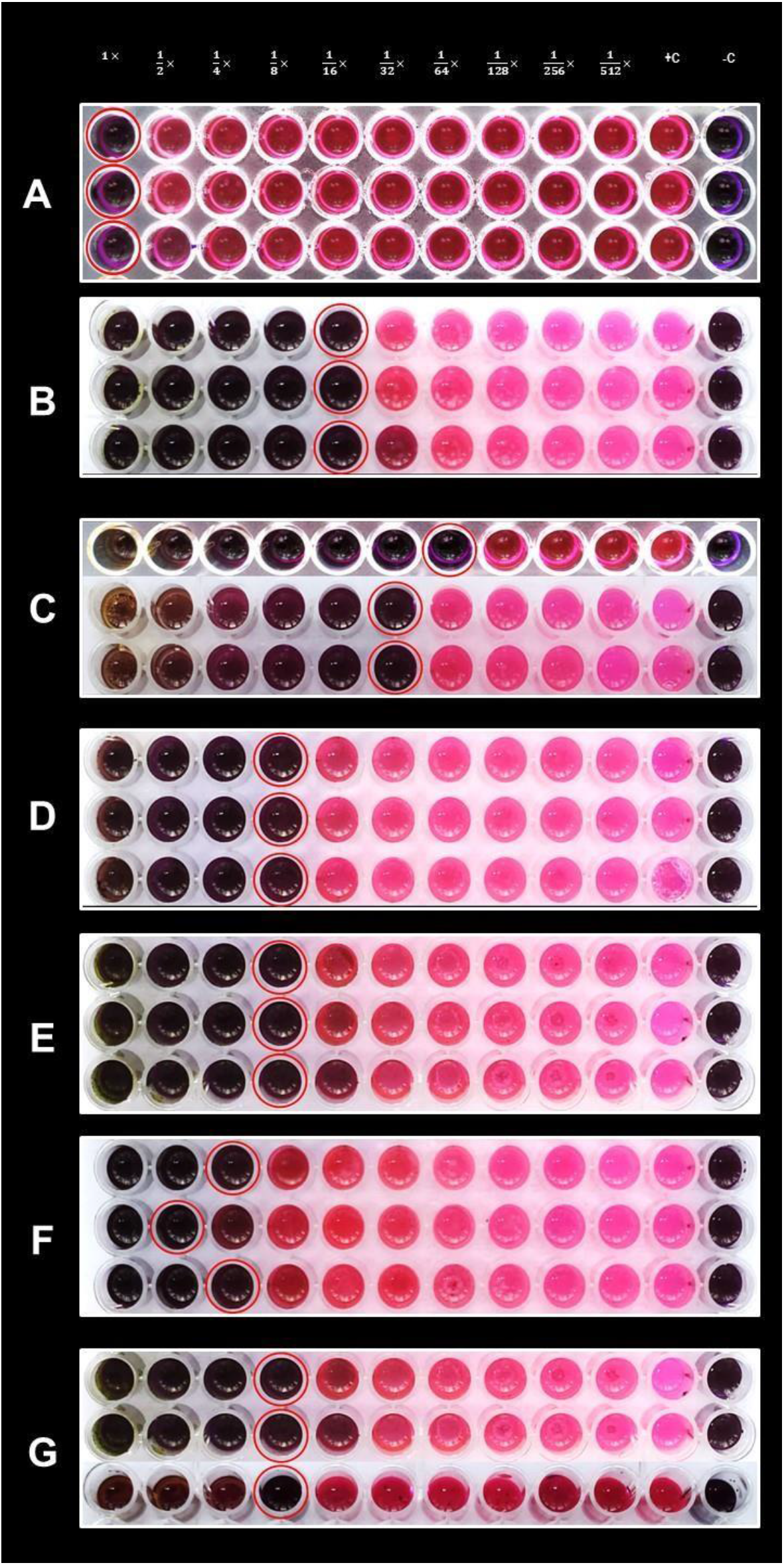
Minimum inhibitory concentrations of the plant extracts and amoxicillin against P. aeruginosa USTCMS 10013 (lowest to highest). *A, Amoxicillin;B, P. betle ethanol extract; C, P. betle methanol extract; D, A. paniculata ethanol extract; E, A. paniculata methanol extract; F, A. paniculata aqueousextract; G, P. betle aqueous extract; +c, positive control; -c, negative control

**Figure 7.**
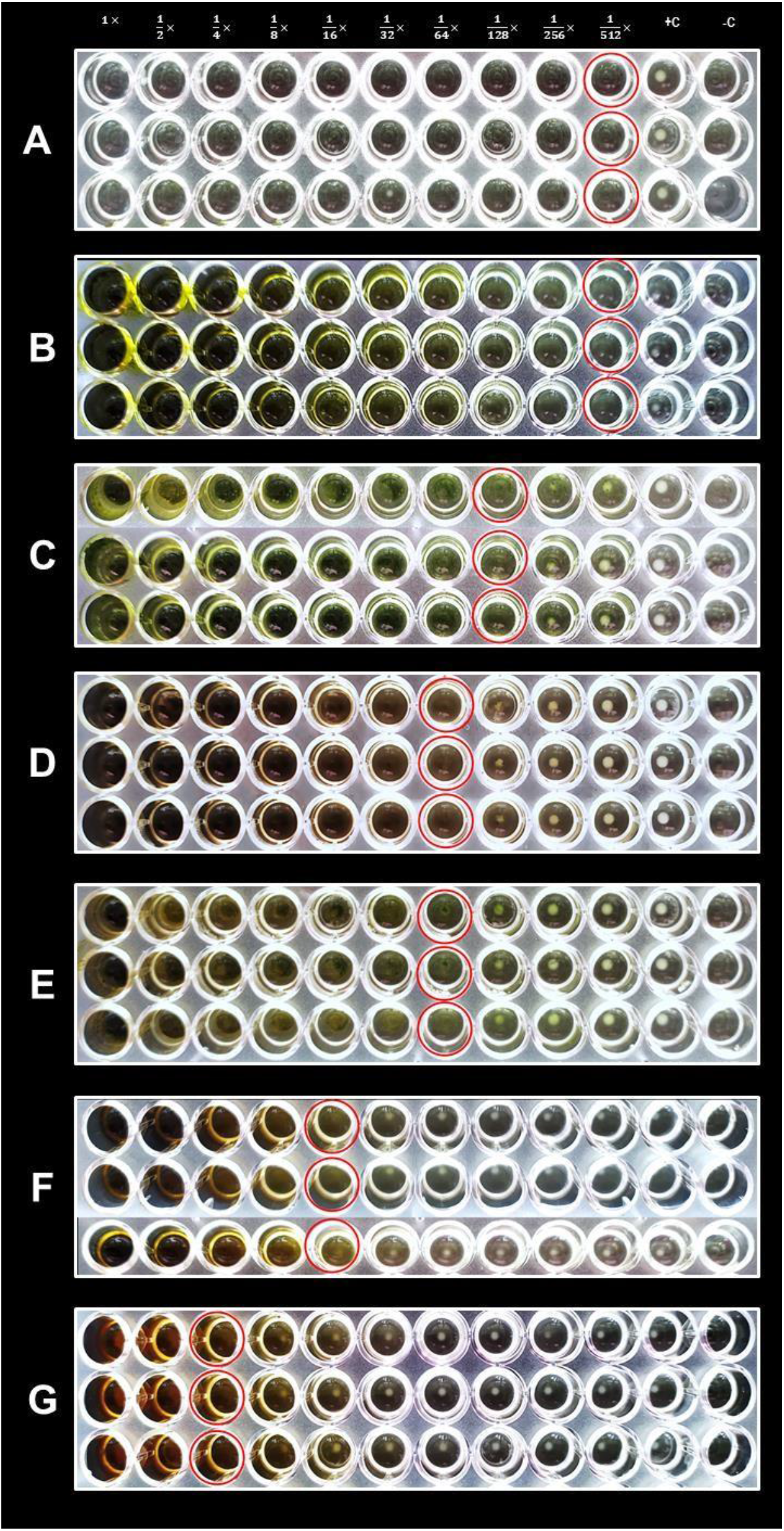
Minimum inhibitory concentrations of the plant extracts and amoxicillin against S. aureus USTCMS 1097 (lowest to highest). *A, Amoxicillin; B, P. betle ethanol extract; C, A. paniculata ethanol extract; D, P. betle methanolextract; E, A. paniculata methanol extract; F, A. paniculata aqueous extract;G, P. betle aqueous extract; +c, positive control; -c, negative control

**Table 3.** Minimum inhibitory concentration of the plant extracts.

| TEST STRAIN | MINIMUM INHIBITORY CONCENTRATION (mg /mL) / EXTRACT |  |  |  |  |  |  |
| --- | --- | --- | --- | --- | --- | --- | --- |
|  | *Andrographis paniculata* |  |  | *Piper betle* |  |  | *Antibiotic |
|  | Methanol | Ethanol | Aqueous | Methanol | Ethanol | Aqueous | Amoxicillin |
| *Escherichia coli* USTCMS 1030 | 231.9 ± 0.0 | 61.2 ± 0.0 | 46.9 ± 9.4 | 35.2 ± 0.0 | 6.5 ± 0.0 | 344.3 ± 0.0 | 0.031 ± 0.005 |
| *Pseudomonas aeruginosa* USTCMS 10013 | 58 ± 0.0 | 30.6 ± 0.0 | 75.1 ± 18.8 | 7.33 ± 1.5 | 3.25 ± 0.0 | 86.1 ± 0.0 | 1 ± 0.0 |
| *Staphylococcus aureus* USTCMS 1097 | 7.24 ± 0.0 | 1.9 ± 0.0 | 23.5 ± 4.7 | 4.4 ± 0.0 | 0.2 ± 0.0 | 122.2 ± 0.0 | <0.002 |

Values are means of triplicate determination (n = 3) ± Standard Error of the Mean (SEM).

It was also observed that both methanol and ethanol extracts of *A. paniculata* and *P. betle* exhibited lower MIC values than those extracted using water. This confirms the findings of Al-Hashimi (2012) that alcoholic extracts have higher antibacterial activity compared to aqueous extracts because higher amounts of phenolic compounds can be extracted when alcohols such as ethanol was used in plant extraction. Another study in the Philippines also detected similar results wherein ethanol extracts of guyabano, ulasimang bato, sambong, and tsaang gubat leaf showed more potent antibacterial activity compared to their aqueous counterparts (Maramba *et al*., 2020).

The ethanol and methanol extracts of *P. betle* leaf displayed lower MICs compared to *A. paniculata*’s ethanol and methanol extracts. Based on MICs, the most potent compound against all the test organisms was the ethanol extract of *P. betle* leaf with MIC values of 6.5, 3.25 and 0.2 mg/mL (Table 3) against *E. coli* USTCMS 1030 (Figure 5), *P. aeruginosa* USTCMS 10013 (Figure 6), and *S. aureus* USTCMS 1097 (Figure 7), respectively. For *A. paniculata* leaf extracts, *E. coli* USTCMS 1030 was best inhibited by its aqueous extract while *S. aureus* USTCMS 1097 and *P. aeruginosa* 10013 were both best inhibited by its ethanol extract (Table 3; Figure 4, 5, & 6).

Results also showed that *P. betle*’s methanol and ethanol extracts have better antibacterial activity compared to *A. paniculata*’s methanol and ethanol extracts against the three test pathogens. Despite not having an inhibition zone when subjected to disk diffusion assay, the ethanol extract of *P. betle* showed the strongest antibacterial activity against all test bacteria as shown in the broth microdilution assay. There were studies that indicate the limitations of the disk diffusion method when it comes to the lack of diffusion of non-polar molecules into the aqueous agar matrix that is used in the assay (Eloff, 2019). Although ethanol is generally a polar solvent, it has a non-polar structure which helps in its ability to also extract non-polar compounds (Permadi *et al*., 2020). It is possible that the ethanol extract of *P. betle* may also contain non-polar molecules which had antimicrobial activity but was not able to diffuse well into the agar. It was only apparent when subjected in broth microdilution.

Meanwhile, the aqueous extract of *A. paniculata* exhibited greater inhibitory activity against *E. coli* USTCMS 1030, *S. aureus* USTCMS 1097 and *P. aeruginosa* USTCMS 10013 than the aqueous extract obtained from *P. betle* (Table 3, Figure 5, 6, & 7). The aqueous extract of *A. paniculata* had MIC values of 23.5, 46.9 and 75.1 mg/mL against *S. aureus*, *E. coli* and *P. aeruginosa*, respectively. On the other hand, the aqueous extract of *P. betle* had MIC values of 86.1, 122.2 and 344.3 mg/mL against *P. aeruginosa*, *S. aureus* and *E. coli*, respectively. These results indicate that the aqueous extract from *A. paniculata* contain a more potent antibacterial compound compared to the aqueous extract from *P. betle*.

The MIC values of the ethanol and methanol extracts of *P. betle* obtained in this study were relatively higher compared to that of the MIC values gathered in the study of Valle *et al*. (2016). Their study demonstrated MIC values of *P. betle*’s ethanol extract of 0.078 to 0.156 mg/mL against strains of methicillin-resistant *S. aureus*. For *E. coli*, they obtained 0.312 mg/mL MIC values for both methanol and ethanol extracts, while for the *P. aeruginosa*, ethanol and methanol extracts had MICs of 0.312 mg/mL and 0.625 mg/mL, respectively. In another study on *A. paniculata*, its methanol extract had an MIC value of 0.031 mg/mL against *E. coli* and *P. aeruginosa* (Wei *et al*., 2011). The results of the present study showed higher inhibitory activity compared to a study by Jamelarin and Balinado (2019) who obtained MIC values of the methanol and ethanol extracts of *P. betle* against *P. aeruginosa* at 150 mg/mL for both extracts.

Results of the present study on the antibacterial property of *A. paniculata* and *P. betle* may not have been as effective as in the study by Valle *et al*. (2016) and Wei *et al*. (2011) due to the modified extraction method. It is also possible that the elevation in the source of the plant may have affected the ability of the plant extracts to inhibit bacterial growth as essential oils and other phytochemical compounds vary depending on the geographical location of the plant (Ermawati *et al*., 2021). Another possibility may also be attributed to the different strains of test bacteria used in various literature compared to this study. The clinical isolates used in this study may be more resistant to various antimicrobials, including plant extracts, compared to other test bacterial strains that are not of clinical nature.

The MIC values of all extracts show that gram-positive bacteria such as *S. aureus* are more susceptible to the antibacterial activity of the plant extracts than gram negative ones such as *E. coli* and *P. aeruginosa*. This demonstrates that gram negative rods are more resistant and need a much higher concentration of antimicrobials present in the plant extracts. This difference in susceptibility may be attributed to the variation in their cell wall structure since gram negative bacteria have an extra outer membrane that consists of lipopolysaccharide and proteins which filter molecules that pass through the cell wall. Gram negative bacteria also have efflux pumps and can more easily acquire resistance genes from mobile genetic elements such as plasmids (Breijyeh *et al*., 2020). It is possible that these characteristics by gram negative bacteria have contributed to their ability to prevent antimicrobial drugs such as antibiotics and phytochemicals from plant extracts from causing damage to the cell thus, making them more resistant compared to gram positive bacteria.

An unusual result was found when resazurin was used for the determination of the MIC of the plant extracts against *S. aureus* USTCMS 1097 in the broth microdilution (Figure 8). Resazurin is a color-based indicator that allows the detection of microbial growth even without the use of a spectrophotometer. A color change from blue-violet to pink indicates that there is a presence of viable cells in the medium (Moondra *et al*., 2018). Resazurin is used in many studies that involve antibacterial determination. In this study, however, it was found out that resazurin can inhibit the growth of *S. aureus* USTCMS 1097. This antimicrobial property of resazurin may be specific only to the strain *S. aureus* USTCMS 1097 since growth is not inhibited in other *S. aureus* strains used by Jamelarin & Balinado (2019) and Teh *et al*. (2017). There are also studies that have been successful in using resazurin and resazurin- based compounds as antimicrobial agents against Francisella tularensis (Schmitt *et al*., 2013) and Neisseria gonorrhoeae (Schmitt *et al*., 2016).

**Figure 8.**
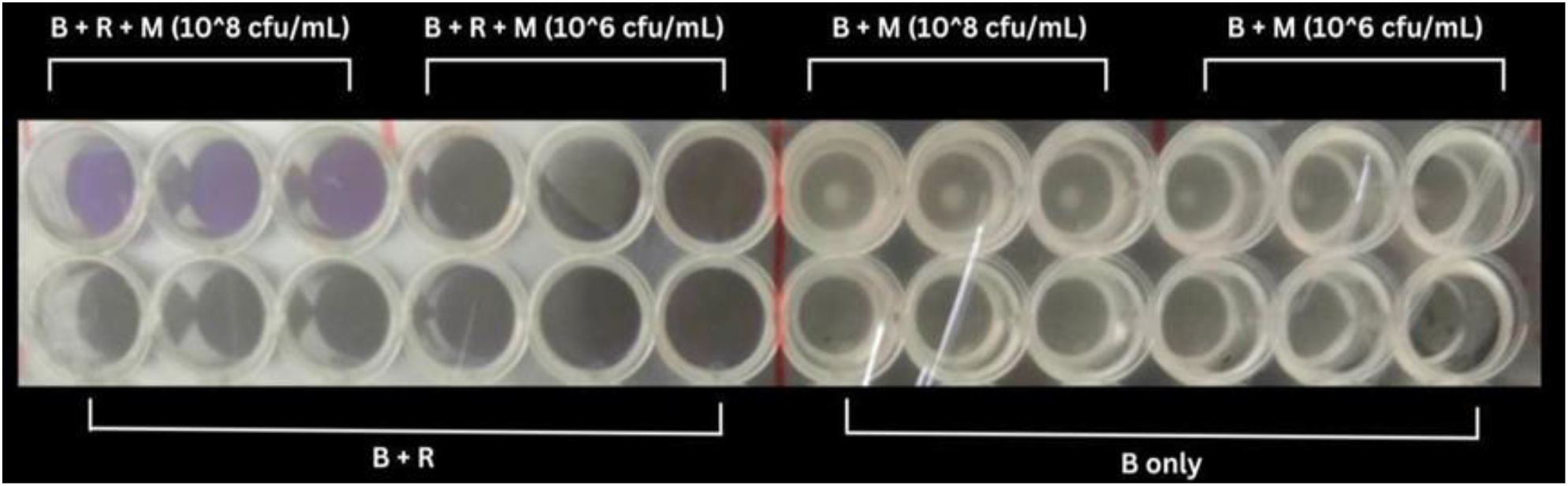
Antibacterial activity of resazurin against *S. aureus* USTCMS 1097 as shown by broth microdilution. \**B, Mueller-Hinton Broth; M, Microbe; and R, Resazurin*

### Interactive Effects of Plant Extracts and Amoxicillin through Disk Diffusion Assay

The individual inhibition zones of the plant extracts and amoxicillin, together with their combined effects are outlined in Tables 4 and 5.

**Table 4.** Interactive effects of A. paniculata extracts and amoxicillin.

| TEST STRAIN | ZONE OF INHIBITION (mm) |  |  |  |  |  |  |  |  |  |  |  |
| --- | --- | --- | --- | --- | --- | --- | --- | --- | --- | --- | --- | --- |
|  | APM | AMX | APM + AMX | INTERACTION | APE | AMX | APE + AMX | INTERACTION | APA | AMX | APA + AMX | INTERACTION |
| *EC* | 0 | 18.2 ± 0.4<sup>a</sup> | 18.2 ± 0.4<sup>a</sup> | IND | 0 | 18 ± 0.0<sup>a</sup> | 17.8 ± 0.9<sup>a</sup> | IND | 0 | 17 ± 0.3<sup>a</sup> | 16.7 ± 0.3<sup>a</sup> | IND |
| *PA* | 0 | 0 | 0 | N | 0 | 0 | 0 | N | 0 | 0 | 0 | N |
| *SA* | 9 ± 0.6<sup>c</sup> | 31.3 ± 0.3<sup>a</sup> | 29.7 ± 0.3<sup>b</sup> | ANT | 10.3 ± 0.4<sup>c</sup> | 31.8 ± 0.2<sup>a</sup> | 29.5 ± 0.5<sup>b</sup> | ANT | 0 | 31.5 ± 0.5<sup>a</sup> | 27.2 ± 0.9<sup>b</sup> | ANT |

Values are means of triplicate determination (n = 3) ± Standard Error of the Mean (SEM). Lowercase letters indicate significant differences (P < 0.05) between different plant extract and amoxicillin combinations against the test strain, where the row was split into three, the first three sub-tables were compared together and the same was done to the succeeding three sub-tables. \*EC, *Escherichia coli* USTCMS 1030; PA, *Pseudomonas aeruginosa* USTCMS 10013; SA, *Staphylococcus aureus* USTCMS 1097; APM, Methanol extract of *A. paniculata*; APE, Ethanol extract of *A. paniculata*; APA, Aqueous extract of *A. paniculata*; AMX, Amoxicillin. Interaction: SYN, Synergistic; IND, Indifferent; ANT, Antagonistic; ADD, Additive; N, No effect.

**Table 5.**
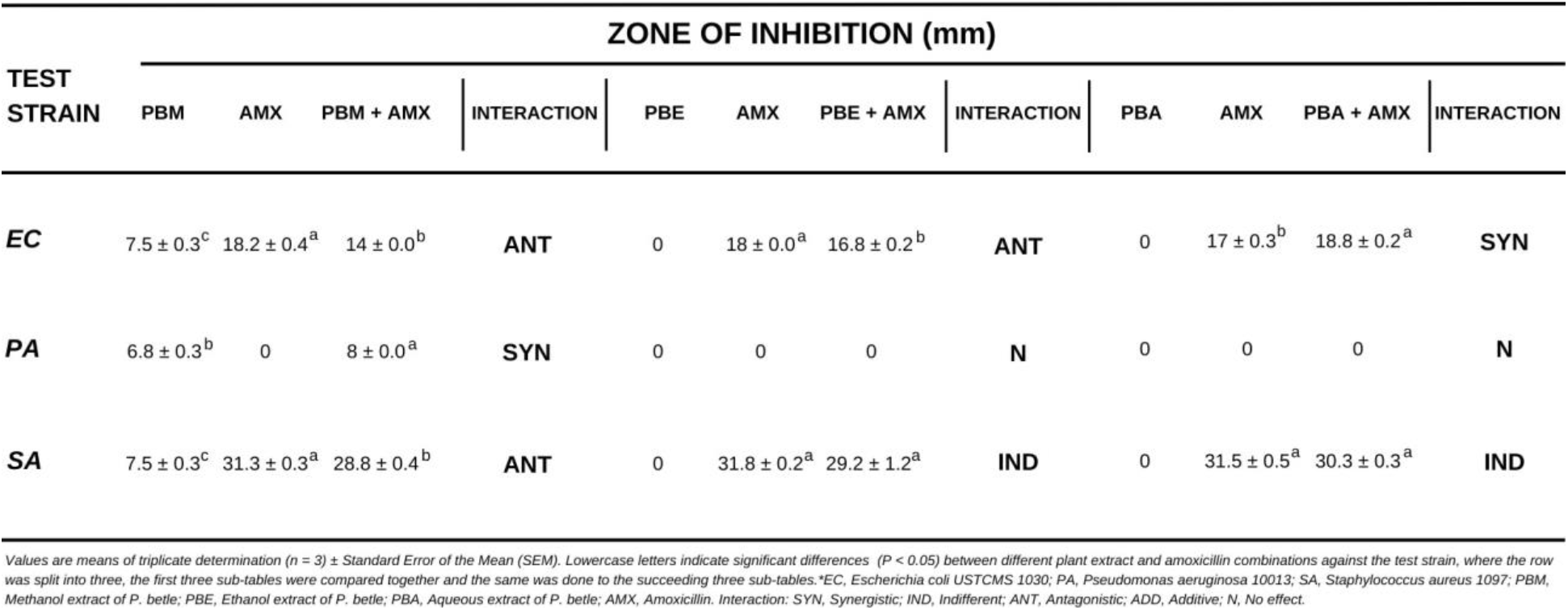
Interactive effects of *P. betle* extracts and amoxicillin.

Against *E. coli* USTCMS 1030, the combination of *A. paniculata* extracts and amoxicillin all resulted in Indifferent interactions while, *P. betle* alcoholic extracts integrated with amoxicillin produced antagonistic interactions. Synergism was evident for aqueous extract with antibiotic against *E. coli* USTCMS 1030 (Figure 9). For *A. paniculata*, the methanol extract with amoxicillin (18.2±0.4 mm), ethanol extract with amoxicillin (17.8±0.9mm), and aqueous extract with amoxicillin (16.7±0.3mm), have no significant differences to their individual inhibition zones; thus, indifferent.

**Figure 9.**
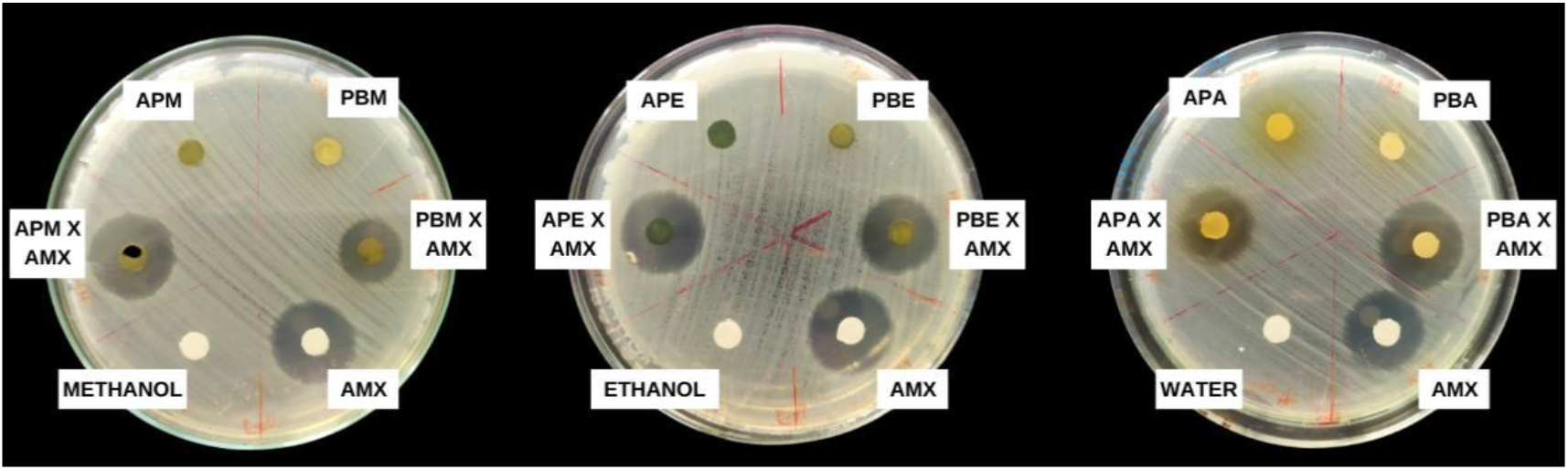
Interactive effects of amoxicillin with A. paniculata and P. betle extracts against E. coli USTCMS 1030. *AMX, Amoxicillin; APM, A. paniculata methanol extract; PBM, P. betle methanol extract; APE, A. paniculata ethanol extract; PBE, P. betle ethanol extract; APA, A. paniculata aqueousextract; PBA, P. betle aqueous extract

The observed antagonism in ethanol extract of *P. betle* with amoxicillin is contradictory with Siahaan’s study in 2016. The *P. betle* ethanol extract combined with amoxicillin resulted in synergism or increased diameters in inhibition zones against *E. coli* which is similar to Sitorus’ (2018) findings. As the results indicate that all *A. paniculata* extracts showed no activity against *E. coli* USTCMS 1030 (Table 4), the activity of plant extract with antibiotics may be solely acted upon by the amoxicillin. The addition of *A. paniculata* extracts does not supplement the existing antibacterial activity of amoxicillin.

On the other hand, two interactive activities were observed in the combination of *P. betle* extracts and amoxicillin against *E. coli* USTCMS 1030; mainly antagonism (ANT) and synergism (SYN). The *P. betle* methanol extract combined with amoxicillin had a zone of inhibition of 14 mm against *E. coli* USTCMS 1030. This measurement is larger than the zone of inhibition of methanol extract alone but is smaller than in amoxicillin alone. This shows antagonism wherein the addition of methanol extract of *P. betle* hindered the effect of the amoxicillin against *E. coli* USTCMS 1030. This finding is similar to the result of Fauziansyah (2019) which showed that the amoxicillin integrated with *P. betle* methanol extract resulted in an antagonistic effect against *E. coli*.

Meanwhile, synergism was observed in combined *P. betle* aqueous extract and amoxicillin against *E. coli* USTCMS 1030. The mean inhibition zone of combined antimicrobial agents has a significant difference to each independent compound (*p=*0.006). Despite showing no inhibitory effects for *P. betle* aqueous extract, when combined with amoxicillin (17±0.3mm), the inhibition zone increased significantly to 18.8±0.2mm.

Against *P. aeruginosa* USTCMS 10013, there were no interactive effects between *A. paniculata* extracts and amoxicillin as shown by the absence of zones of inhibition when used individually or combined (Table 4; Figure 10). The same result was obtained when tested with *P. betle*’s ethanol and aqueous extracts. Only the *P. betle* methanol extract showed inhibitory activity against *P. aeruginosa* USTCMS 10013. This activity was significantly increased when this plant extract was combined with amoxicillin (Table 5; Figure 10). This result reveals synergism between methanol extract of *P. betle* and the antibiotic amoxicillin in inhibiting *P. aeruginosa* USTCMS 10013. This combination of antimicrobial agents may be a potential candidate for treating strains of amoxicillin-resistant *P. aeruginosa*.

**Figure 10.**
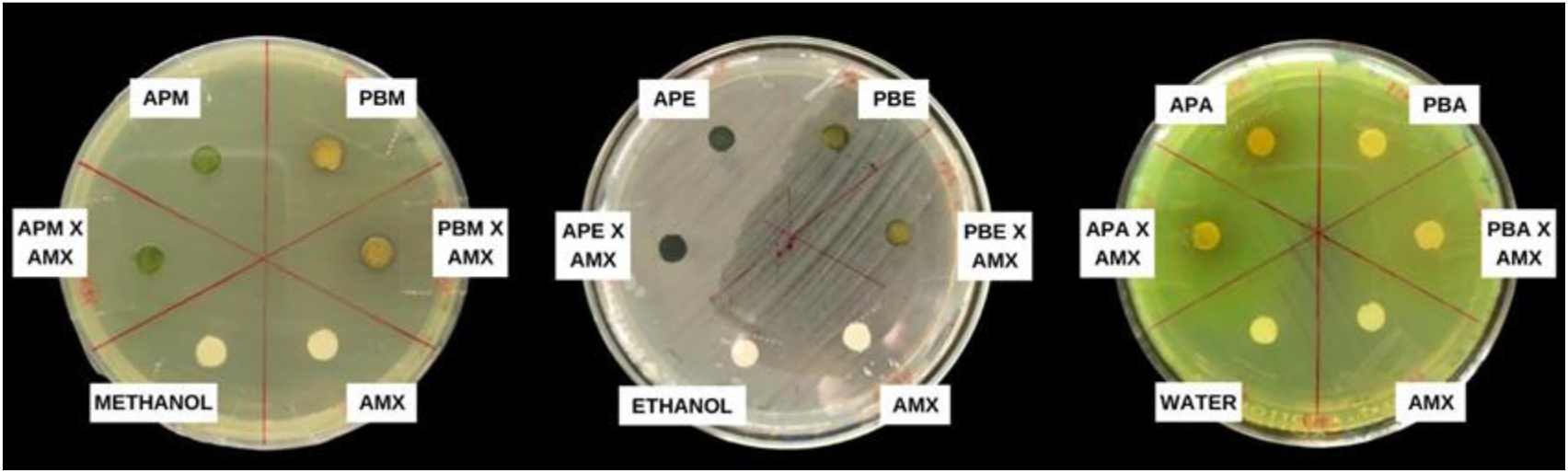
Interactive effects of amoxicillin with A. paniculata and P. betle extracts against P. aeruginosa USTCMS 10013. *AMX, Amoxicillin; APM, A. paniculata methanol extract; PBM, P. betle methanol extract; APE, A. paniculata ethanol extract; PBE, P. betle ethanol extract; APA, A. paniculata aqueous extract; PBA, P. betle aqueous extract

**Figure 11.**
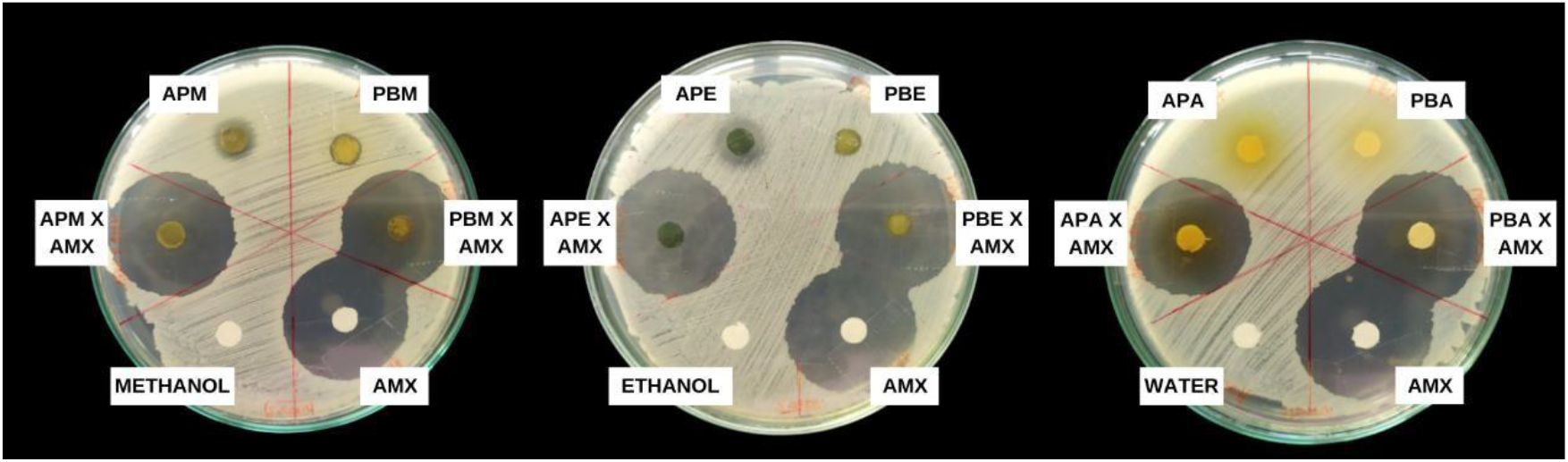
Interactive effects of amoxicillin with A. paniculata and P. betle extracts against S. aureus USTCMS 1097. *AMX, Amoxicillin; APM, A. paniculata methanol extract; PBM, P. betle methanol extract; APE, A. paniculata ethanol extract; PBE, P. betle ethanol extract; APA, A. paniculata aqueous extract; PBA, P. betle aqueous extract

Phytochemicals with antimicrobial compounds such as terpenoids, saponins, tannins, flavones (Malkhan *et al.,* 2012), and phenolic constituents such as eugenol and hydroxychavicol (Nagabhushan *et al.,* 1989) can be extracted using methanol. Furthermore, methanol extract of *P. betle* contains hydroxychavicol (Nguyen *et al.,* 2020) which is a genotoxic compound that can induce cell impairment and disrupt cell division of bacterial cells. These compounds could be responsible for the synergistic activity of the methanol extract of *P. betle* to amoxicillin against *P. aeruginosa* USTCMS 10013.

Against *S. aureus* USTCMS 1097, antagonism was observed on the interactive effects of *A. paniculata* extracts and amoxicillin as well as in the combination of *P. betle* methanol extract and amoxicillin. Indifference was observed in *P. betle* ethanol and aqueous extracts combined with amoxicillin. Results indicate that all the *A. paniculata* extracts and methanol extract of *P. betle* do not give supplementary effect, but have instead weakened the antimicrobial activity of amoxicillin in inhibiting *S. aureus* USTCMS 1097.

In other studies, ethanol extract of *P. betle* combined with amoxicillin showed synergistic effect against *S. aureus* (Siahaan, 2016; Sitorus, 2018). This is not the same as the effect found in the present study. However, Fauziansyah (2019) mentioned that the amoxicillin integrated with *P. betle* methanol extract resulted in an antagonistic effect against *S. aureus*, a finding that is similar to this study.

### Interactive Effects of Plant Extract and Amoxicillin through Checkerboard Assay

Checkerboard assay was used to quantitatively evaluate the synergistic effect of plant extracts with amoxicillin. Only the combinations that showed synergism using disk diffusion method were subjected in this assay. The FICI values of the interactive effects of the subjected plant extracts with amoxicillin is shown in Table 6.

**Table 6.** Interaction between P. betle extracts with amoxicillin expressed as FICI.

| TEST STRAIN | ANTIMICROBIAL AGENTS<br>IN COMBINATION | FRACTIONAL INHIBITORY CONCENTRATION INDEX (FICI) |  |  |  |
| --- | --- | --- | --- | --- | --- |
|  |  | FIC ant | FIC ext | Σ FIC | INTERACTION |
| *Escherichia coli*<br>USTCMS 1030 | AMX + PBA | 0.33 | 0.33 | 0.66 | ADDITIVE |
| *Pseudomonas aeruginosa*<br>USTCMS 10013 | AMX + PBM | 0.01 | 0.13 | 0.14 | SYNERGISTIC |

Synergistic (FICI ≤ 0.5) if the effect of the two agents was stronger than the sum of effects of the individual agents; Additive (FICI > 0.5-1) if their joint effect was identical to the overall effects of the individual agents; Antagonistic (FICI > 4) if the interaction exhibits a weaker result than the sum of effects of the individual agents, or weaker than the effect of either individual agent; and Indifference (FICI > 1-4) if the result of their interaction was equal to the overall effect of either individual agent. \*AMX, Amoxicillin; PBA, Aqueous extract of *P. betle*; PBM, Methanol extract of *P. betle*.

The results of the combined effect of amoxicillin and the aqueous extract of *P. betle* in the checkerboard assay did not acquire the same interactive result as in the preliminary disk diffusion assay. The resulting interactive effect as expressed in FICI is 0.66 and is interpreted as only additive. This indicates that the activity of the amoxicillin was not enhanced or hindered by the plant extract, but that their individual effects have just added together (Figure 12). *E. coli* has long been known to be a causative agent of many diseases, and studies show its role in lower respiratory tract infections such as pneumonia (Edwards *et al.,* 2019). Its resistance to antimicrobials has been on the rise, and so novel approaches of fighting these bacteria such as employing combination therapy, examples of which is the merging of antibiotics with chemicals from plant extracts, are now needed more than ever.

**Figure 12.**
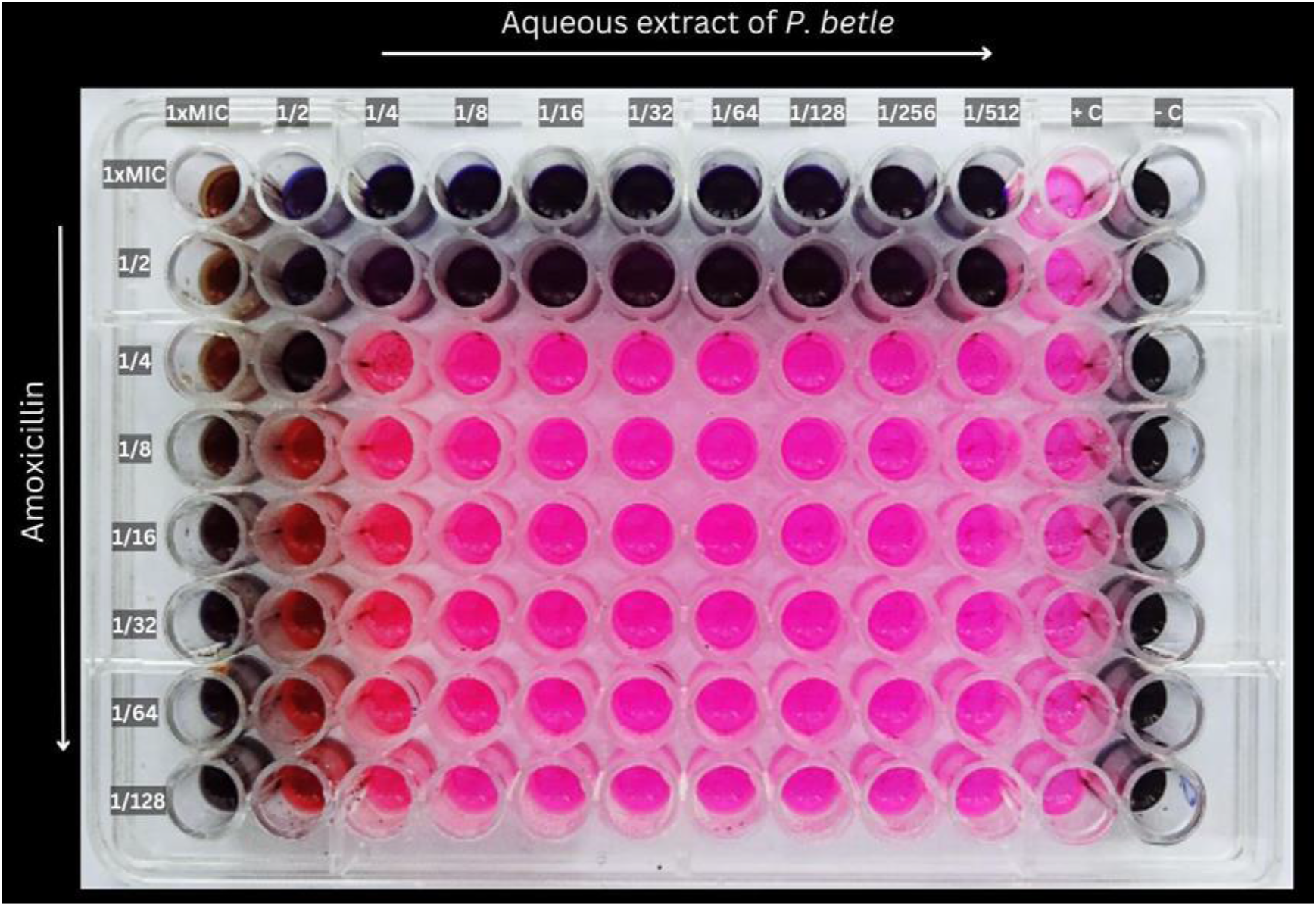
Checkerboard Assay of the aqueous extract of *P. betle* and amoxicillin against *E. coli* USTCMS 1030*. +c, positive control; -c, negative control*

FICI results from the checkerboard assay (0.33, Synergistic) confirmed the synergism between methanol extract of *P. betle* and amoxicillin against *P. aeruginosa* USTCMS 10013. This plant extract had effectively strengthened the antibacterial activity of the amoxicillin by 128-fold (Figure 13). This finding is the first observation of the synergistic effect of the methanol extract of *P. betle* combined with amoxicillin against *P. aeruginosa*. Other studies only recorded synergistic effects of the ethanol and ethyl acetate extract of *P. betle* with chloramphenicol and streptomycin against some strains of *S. aureus* and *P. aeruginosa* (Taukoorah *et al.,* 2016). *P. aeruginosa* is one of the most often isolated pathogens in the respiratory system of people with LRTIs (Gellatly & Hancock, 2013). They are also known to be very resistant to beta- lactam antibiotics such as amoxicillin (Glen & Lamont, 2021). This synergism of amoxicillin with *P. betle* may be explored for the treatment of respiratory infections that are caused by *P. aeruginosa*.

**Figure 13.**
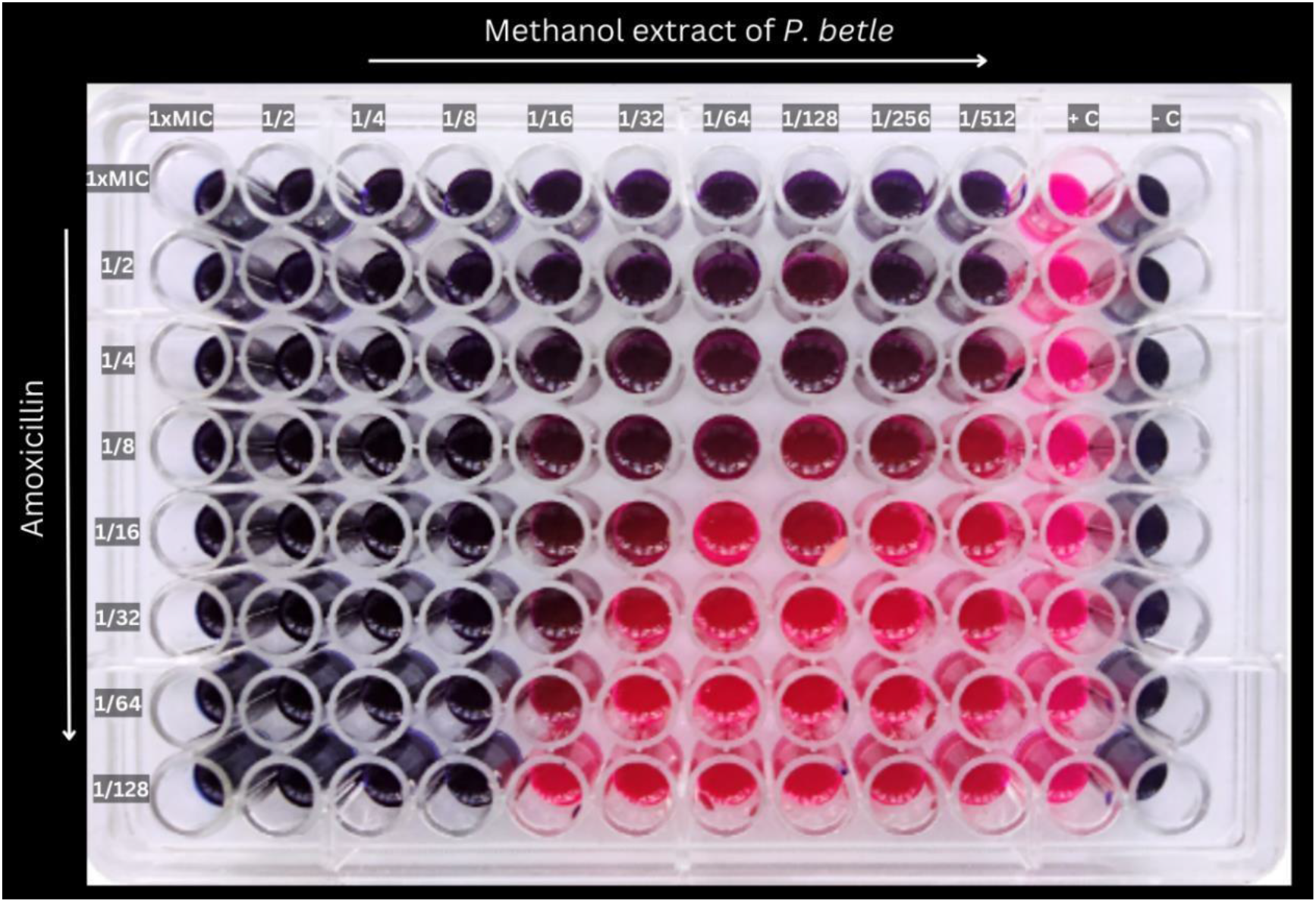
Checkerboard assay of the methanol extract of *P. betle* and amoxicillin against *P. aeruginosa* USTCMS 10013. *+c, positive control; -c, negative control*

*P. betle* contains alkaloids, phenols, tannins and flavonoids which all have potential antibacterial activity (Syahidah *et al.,* 2017). Solvents such as water, ethanol, and methanol may extract different active components that have proven antimicrobial activities. Plant extraction using water as solvent, yields terpenoids, tannins, and saponins. Ethanol yields compounds like alkaloids, tannins, terpenoids, and flavonoids. Terpenoids, saponins, tannins, and flavonoids are extracted using methanol (Malkhan *et al.,* 2012).

Moreover, a study by Rekha *et al*. (2014) indicated that Chavibetol, which is a phenol, is the main chemical component of a *P. betle* leaf. Hydroxychavicol is another phenol with antimicrobial activity that was reported to be extracted by methanol in *P. betle* leaves (Nguyen *et al*., 2020). Phenols have an ability to impair some essential enzymes as it has high protein binding affinity to microbial protein complexes (Miklasińska-Majdanik *et al*., 2018; Aldulaimi, 2017). Flavonoids have abilities such as being able to impede porin function, affect cytoplasmic membrane and inhibit nuclear synthesis (Xie *et al*., 2014) whereas tannins and alkaloids interfere with cell metabolism and form complexes with the bacterial cell wall (Kaczmarek, 2020; Othman *et al*., 2019). The synergistic interaction between the methanol and aqueous extract of *P. betle* with amoxicillin may possibly be attributed to the activity of the bioactive components of the plant such as phenols which disrupt the enzymes responsible for the amoxicillin resistance in the *E. coli* and *P. aeruginosa*. These resistance mechanisms against antibiotics like amoxicillin can be caused by beta- lactamases, increase in efflux pumps, and reduction of antibiotic uptake by means of porins (Glen & Lamont, 2021). The bioactive components of the plant may have acted as Resistance Modifying agents as it disrupted the mechanisms causing the antibiotic resistance, thereby facilitating and promoting the effect of the antibiotic inside the cell of the bacteria.

Combination therapy, which is the use of two or more antimicrobial agents simultaneously, which in this case is between a plant extract and an antibiotic, could be a promising approach towards developing novel antimicrobial drugs or resistance modifying agents that will regain the effect of overused antibiotics. This approach can also potentially be highly effective since it will be difficult for a bacterial cell to mutate resistance mechanisms that will make it resistant to numerous drugs with different antibacterial activities (Talaro, 2008).

## MATERIALS AND METHODS

### Collection of Samples and Pre-extraction Preparation

*paniculata* and *P. betle* plants to be subjected to extraction were collected in Cavite. Whole live plant of *A. paniculata* was collected from Naic, Cavite. The *P. betle* plant was obtained from Tanza, Cavite. The section of each whole plant was submitted to and authenticated by the Jose Vera Santos Memorial Herbarium of the Institute of Biology, College of Science, University of the Philippines – Diliman. After confirmation, the leaves of each plant were collected, washed using tap water, then by distilled water. Afterwards, the leaves were air dried for three to four days. The leaves were grounded into powdered form using an electric blender. The dried plant materials were weighed and then subjected into maceration using three different solvents: analytical grade of methanol and ethanol and distilled water.

### Crude Leaf Extraction (Double Maceration)

The method of double maceration was adapted from Balakrishna’s group (2016) and Singh (2008) with slight modifications. Initially, 20g plant samples were macerated in 100 ml solvent for 72 hours with periodic churning. The crude extract was obtained by filtration using a clean muslin cloth. The crude extract was placed inside a clean, secured container and was stored at room temperature. Then the same plant material was added with another 100 ml solvent for further extraction of active compounds. This second maceration was done for 24 hours. The same procedure on the first maceration was followed. The crude extracts obtained were mixed with extract from the first maceration, and were allowed to stand for two more weeks. The solution was then filtered using Whatman Filter Paper No. 1 and the filtrate was placed in a beaker. The filtrates were then transferred to conical flasks, sealed and stored at room temperature.

### Evaporation of Solvent

The methanol and ethanol extracts of the plant samples were further processed using a Rotary Evaporator. This process evaporated the solvent to obtain a pure concentration of the plant extract. As for the aqueous extract of the plant samples, a freeze dryer was utilized to remove the water, leaving a concentrated plant extract.

The rotary evaporator was used to further evaporate remaining alcohols and discriminated it from the plant extract. The filtrate was placed into a round-bottom flask and firmly attached to the Rotary Evaporator. The Rotary Evaporator was manipulated according to the Standard Operating Procedure (SOP). The rotation per minute (rpm) was set into 25 rpm. The round- bottom flask with filtrate was lowered into the heating bath. The flask containing the extract was then removed from the heating bath after the liquid was partially evaporated and left a viscous distinctive adhesion of plant material at the bottom. The small amount of the final product was contained in a small sealed semi-ventilated amber vial. Lastly, the vial was placed in the fume hood overnight for further evaporation. The end products were refrigerated at 4°C until further utilization.

Twenty mL of the aqueous plant extracts were placed in a 50-mL falcon tube. Samples were frozen at a temperature of -86°C which were then subjected to freeze drying. The Falcon tubes screw caps were replaced with perforated parafilm. Then, each identical extract was placed in a separated fast-freeze flask that was attached to the valve ports. Freeze drying was done continuously for three days. The dried products were collected, weighed, kept in a sealed amber vial, and refrigerated until further use.

The extraction yield of each plant was calculated using the following equation (Felhi *et al.,* 2017).

**Extraction Yield (%) = (X1 ∗ 100)/X0**

Where X1 refers to the weight of extract after evaporation of solvent and X0 refers to the dry weight of the plant powder before extraction.

### Preparation of Plant Extract Stock Solution

Each concentrated plant extract was subjected to their respective solvents (water, ethanol, and methanol). A stock solution for each preparation was constructed by dissolving the entire yield of extract with 6 mL of their respective solvents. The reconstituted plant extracts were stored in sealed amber vials which were refrigerated until further use. Sterility of extracts was ensured by plating them on Nutrient Agar (NA) before the experiment was performed.

### Preparation of Bacterial Culture

The test microorganisms used were clinical isolates of *Staphylococcus aureus* USTCMS 1097*, Escherichia coli* USTCMS 1030, and *Pseudomonas aeruginosa* USTCMS 10013 obtained from the University of Santo Tomas Collection of Microbial Strains (USTCMS). These were cultivated and maintained in Nutrient Agar (NA) plate, slants and Nutrient Broth (NB) at 4°C.

All the culture media used in the maintenance and assay of these bacterial cultures were prepared and sterilized according to standard procedures and manufacturer’s instruction.

### Preparation of Antibiotics

Amoxicillin in capsules (AMBIMOX) were used. Stock solution (1 mg/mL) of the antibiotic was prepared by weighing and dissolving 5 mg of amoxicillin powder into a 5 mL sterile H2O. The amoxicillin solution was further homogenized using a vortex mixer. It was then filter-sterilized and stored in a refrigerator.

### Antimicrobial Susceptibility Testing

Disk Diffusion Assay (Penecilla, 2011) was used for the antimicrobial susceptibility testing of the three pathogens to the plant extracts. Each bacterial culture was grown in NA at 37°C overnight. Cells were suspended in normal saline solution. The turbidity was adjusted to equal the turbidity of the 0.5 McFarland Standard to get an inoculum of approximately 1.5x108 CFU/mL. A sterile cotton swab was dipped into the bacterial suspension and then swabbed and spread evenly over the entire agar plate surface. The filter paper disks were impregnated with the 20 µL plant extracts and were allowed to dry for 15 to 30 minutes prior to placing on top of the Mueller-Hinton Agar with the test organism. Filter paper disks with only the solvent were used as the negative control, while filter paper disks that are impregnated with antibiotic solution were used as the positive control. The agar plates with the disks were allowed to stand for an hour at room temperature. The plates were then incubated at 37°C for. After 16 to 24 hours, zone of inhibition was measured. An inhibition zone of 14 mm or greater (including diameter of the disk, 6 mm) was considered as high antimicrobial activity. All treatments were done in triplicates to ensure the reliability of the data.

The MIC of plant extracts and antibiotics was determined using broth microdilution by two-fold serial dilution. The 96-well plates were prepared by dispensing 100 μl of Mueller–Hinton broth into each well. A 100 μl from the stock solution of *A. paniculata* and *P. betle* extracts were added into the first row of the plate. Then, twofold serial dilution was performed using a micropipette in order to reduce the extract concentration until the 10th well. The colonies of the test organism that was subcultured on a NA plate at 37°C overnight was diluted in a saline solution. The turbidity was adjusted to equal the turbidity of the 0.5 McFarland Standard to get an inoculum of approximately 1.5x108 CFU/mL. The adjusted inoculum was further diluted by transferring its 100 µL to a 9.9 mL saline solution to get an inoculum of approximately 1.5 x106 CFU/mL. Twenty μl of inocula were added to each well including the 11th well. Ten μl of membrane-filtered resazurin was then added to each well. The 11th well served as the positive control. Whereas, the 12th well served as the negative control which contained Mueller-Hinton Broth (MHB) only. The plates were incubated under 37°C for 16 to 24 h. The use of resazurin was only applied to MIC tests for *E. coli* USTCMS 1030 and *P. aeruginosa* USTCMS 10013. In the case of the *S. aureus* USTCMS 1097, no resazurin indicator was added. Instead, only the wells with bacterial pellets were considered as a definite growth. The amount of growth in each well was compared to the positive control wherein the MIC was the lowest concentration of the extract which showed inhibition to the bacteria.

### Evaluation of Interactive Effects of Plant Extract with Antibiotics

Disk diffusion method described above, was used to determine the interactive effects of the plant extracts (Penecilla, 2011). The filter paper disks were impregnated with 10µL of amoxicillin (1mg/mL) and were allowed to dry for 10-15 minutes. Then, 10µL of plant extracts of known concentration were dropped to the disks before placing them on the agar plate. Filter paper disks with only the solvent were used as the negative control, while filter paper disks that were impregnated with antibiotic solution were used as the positive control. The agar plates were allowed to settle for at least 30 minutes at room temperature before incubating at 37°C for 16-20 hours. After incubation, the diameter of the zones of inhibition was measured using a ruler in millimeters. The interactions of the plant extract and antibiotic were interpreted as follows:

**Synergism -** if the diameter of the zone of inhibition of the two agents is significantly larger than the sum of effects of the individual agents.

**Additive -** if the diameter of the zone of inhibition of the two agents is equal to the sum of effects of the individual agents.

**Indifferent -** if the diameter of the zone of inhibition of the two agents is equal to the overall effect of either individual agent.

**Antagonistic -** if the diameter of the zone of inhibition of the two agents is significantly smaller than the sum of effects of the individual agents.

The checkerboard method of Stefanovic (2018) was used to quantitatively determine the interaction between the antibiotic and the plant extracts from the disk diffusion assay with some modifications. Fifty (50) μl of Mueller-Hinton broth were prepared and distributed into each well of the 96-well microtiter plate. A 50 μl of the MIC of the plant extract was added in the first row of the plates which underwent two-fold serial dilutions to obtain different concentrations of the plant extract. Then, 50 μl of amoxicillin in different concentrations (1xMIC, 1/2xMIC, 1/4xMIC, 1/8xMIC, 1/16xMIC, 1/32xMIC, 1/64xMIC, and 1/128xMIC) were added. Twenty μl of the bacterial suspension at approximately 1.5 x 106 CFU/mL was added to the wells. Finally, 10 μl of resazurin was added to each well. The 11th column which contained the inoculum and the broth served as the positive control. Whereas, the 12th column, which contained only the Mueller Hinton broth, served as the negative control. The 96-well plates were incubated at 37°C for 16-20 hours.

The fractional inhibitory concentration index (FICI) was calculated using the formula:

**FICI = FIC of Antibiotic + FIC of Plant Extract**

Wherein:

**FIC of Antibiotic** = MIC of Antibiotic in Combination / MIC of Antibiotic Alone

**FIC of Plant Extract** = MIC of Plant Extract in Combination / MIC of Plant Extract Alone

The results of the interaction were categorized as synergistic, additive, antagonistic, and indifferent effects as follows.

**Synergistic (FICI ≤ 0.5),** if the effect of the two agents is stronger than the sum of effects of the individual agents;

**Additive (FICI = 0.5-1)**, if their joint effect is identical to the overall effects of the individual agents;

**Antagonistic (FICI > 4)**, if the interaction exhibits a weaker result than the sum of effects of the individual agents, or weaker than the effect of either individual agent;

**Indifference (FICI = 1-4),** if the result of their interaction is equal to the overall effect of either individual agent (Stefanović, 2012).

### Data Processing and Analysis

The data regarding the inhibition zone diameters obtained from the study were represented by means of three replicates ± Standard Error of the Mean (SEM). Data were analyzed by employing one-way analysis of variance (ANOVA), and the mean comparisons were executed by Tukey’s post hoc multiple comparison test using BrightStat software. Differences between means were considered significant at p- value < 0.05.

## REFERENCES

Abreu, A. C., McBain, A. J., & Simões, M. (2012). Plants as sources of new antimicrobials and resistance-modifying agents. Natural Product Reports, 29(9), 1007. https://doi:10.1039/c2np20035j.

Aldulaimi, O. (2017). General overview of phenolics from plant to laboratory, good antibacterials or not. Pharmacognosy Reviews, 11(22), 123. https://doi.org/10.4103/phrev.phrev_43_16.

Al-Hashimi, A. G. (2012). Antioxidant and antibacterial activities of *Hibiscus sabdariffa* L. extracts. African Journal of Food Science 6, 506–511.

Ali, I., Khan, F. G., Suri, K. A., Gupta, B. D., Satti, N. K., Dutt, P., Afrin, F., Qazi, G. N., & Khan, I. A. (2010). In vitro antifungal activity of hydroxychavicol isolated from *Piper betle L*. Annals of Clinical Microbiology and Antimicrobials, 9(1), 7. https://doi.org/10.1186/1476-0711-9-7.

Anderson, L. A. (2021, September 17). Antibiotics Guide. Drugs.com. https://www.drugs.com/article/antibiotics.html.

Azahar, N. I., Mokhtar, N. M., Mahmood, S., Abd Aziz, M. A., & Arifin, M. A. (2023). Evaluation of *Piper betle* L. extracts and its antivirulence activity towards *P. aeruginosa*. Jurnal Teknologi, 85(1). 133–140.

Balakrishna, T., Vidyadhara, S., Sasidhar, R. L. C., Ruchitha, B., & Prathyusha, E. V. (2016). A review on extraction techniques. Indo American Journal of Pharmaceutical Sciences, 3(8), 880–891.

Breijyeh, Z., Jubeh, B., & Karaman, R. (2020). Resistance of gram-negative bacteria to current antibacterial agents and approaches to resolve it. Molecules, 25(6), 1340. https://doi.org/10.3390/molecules25061340.

Cardoso, T. C., Lopes, L. M., & Carneiro, A. H. (2007). A case-control study on risk factors for early-onset respiratory tract infection in patients admitted in ICU. BMC Pulmonary Medicine, 7(1). https://doi.org/10.1186/1471-2466-7-12

Dugassa, J., & Shukuri, N. (2017). Review on antibiotic resistance and its mechanism of development. Journal of Health, Medicine and Nursing, 1(3),1–17.

Edwards, B. D., Somayaji, R., Greysson-Wong, J., Izydorczyk, C., Waddell, B., Storey, D. G., Rabin, H. R., Surette, M. G., & Parkins, M. D. (2019). Clinical outcomes associated with *Escherichia coli* infections in adults with cystic fibrosis: a cohort study. Open Forum Infectious Diseases, 7(1), 1–6. https://doi.org/10.1093/ofid/ofz476.

Eloff, J. N. (2019). Avoiding pitfalls in determining antimicrobial activity of plant extracts and publishing the results. BMC Complementary and Alternative Medicine, 19(1), 1–8. https://doi.org/10.1186/s12906-019-2519-3.

Ermawati, F., Sari, R., Putri, N., Rohmawati, L., Kusumawati, D., Munasir, & Supardi, Z. (2021). Antimicrobial activity analysis *of Piper betle* Linn leaves extract from Nganjuk, Sidoarjo and Batu against *Escherichia coli*, *Salmonella* sp., *Staphylococcus aureus* and *Pseudomonas aeruginosa*. Journal of Physics: Conference Series, 1951(1), 1–10. https://doi.org/10.1088/1742-6596/1951/1/012004.

Fair, R. J., & Tor, Y. (2014). Antibiotics and bacterial resistance in the 21st century. Perspectives in Medicinal Chemistry, 6, PMC-S14459.

Fauziansyah, R. M. (2019). Efek kombinasi ekstrak metanol daun sirih (*Piper betle* Linn.) Dengan antibiotik amoxicillin, chloramphenicol dan cotrimixazole terhadap daya hambat pertumbuhan bakteri *S. aureus* dan *E. coli* [Effect of Combination of Betel Leaf Methanol Extract (*Piper betle* Linn.) with Antibiotics Amoxicillin, Chloramphenicol and Cotrimixazole on the Inhibition of *S. aureus* and *E. coli* bacteria growth. Journal of Community Medicine]. Jurnal Kedokteran Komunitas, 6(3).

Fletcher, J. (2019, February 11). Lower respiratory tract infections: What to know. Medicalnewstoday.com; Medical News Today. https://www.medicalnewstoday.com/articles/324413.

Foo, L. W., Salleh, E., & Mamat, S. N. H. (2015). Extraction and qualitative analysisof *Piper betle* leaves for antimicrobial activities. International Journal of Engineering Technology Science and Research, 2(2), 1–8.

Forum of International Respiratory Societies. (2017). The global impact of respiratory disease. Second Edition. Sheffield, European Respiratory Society. [9781849840880].

Gellatly, S. L., & Hancock, R. E. W. (2013). *Pseudomonas aeruginosa*: new insights into pathogenesis and host defenses. Pathogens and Disease, 67(3), 159–173. https://doi.org/10.1111/2049-632x.12033.

Glen, K. A., & Lamont, I. L. (2021). Β-lactam resistance in *Pseudomonas aeruginosa*: current status, future prospects. Pathogens, 10(12), 1638. https://doi.org/10.3390/pathogens10121638.

Jamelarin, E. M., & Balinado, L. O. (2019). Evaluation of antibacterial activity of crude aqueous, ethanol and methanol leaf extracts of P*iper retrofractum* vahl.and *Piper betle* L. Asian Journal of Biological and Life Sciences, 8(2), 63–67. https://doi.org/10.5530/ajbls.2019.8.11.

Kaczmarek, B. (2020). Tannic acid with antiviral and antibacterial activity as a promising component of biomaterials—a minireview. Materials, 13(14), 3224. https://doi.org/10.3390/ma13143224.

Kraus, E. M., Pelzl, S., Szecsenyi, J., & Laux, G. (2017). Antibiotic prescribing for acute lower respiratory tract infections (LRTI) – guideline adherence in the German primary care setting: An analysis of routine data. PLOS ONE, 12(3), e0174584. https://doi.org/10.1371/journal.pone.0174584.

Knight, G.M., Glover, R.E., Mcquaid, C.F., Olaru, I.D., Gallandat, K., Leclerc, Q.J., Fuller, N.M., Willcocks, S.J. & Hasan, R., van Kleef, E., & Chandler, C.I. (2021). Antimicrobial resistance and COVID-19: intersections and implications. eLife Sciences. 10.10.7554/eLife.64139.

Kumoro, A. C., Hasan, M., & Singh, H. (2009). Effects of solvent properties on the Soxhlet extraction of diterpenoid lactones from *Andrographis paniculata* leaves. Science Asia, 35(1), 306–309.

Malkhan S, Shahid A, Kangabam MS (2012). Efficacy of plant extracts in plant disease management. International Research Journal of Microbiology. pp. 4–23.

Maramba-Lazarte, C. C., Cavinta, L. L., & Sara, M. C. L. (2020). Antibacterial activity of guyabano, ulasimang bato, sambong, and tsaang gubat leaf extracts against common drug-resistant bacteria. Acta Medica Philippina, 54(1), 17–21. https://doi.org/10.47895/amp.v54i1.1087.

Miklasińska-Majdanik, M., Kępa, M., Wojtyczka, R., Idzik, D., & Wąsik, T.J. (2018). Phenolic compounds diminish antibiotic resistance of *Staphylococcus aureus* clinical strains. International Journal of Environmental Research and Public Health, 15(10), 2321. https://doi.org/10.3390/ijerph15102321.

Moondra, S., Raval, N., Kuche, K., Maheshwari, R., Tekade, M., & Tekade, R. K. (2018). Sterilization of pharmaceuticals. Dosage Form Design Parameters, 467–519. doi:10.1016/b978-0-12-814421-3.00014-2.

Nagabhushan, M., Amonkar, A. J., Nair, U. J., D’Souza, A. V., & Bhide, S. V. (1989). Hydroxychavicol: a new anti-nitrosating phenolic compound from betel leaf. Mutagenesis, 4(3), 200–204.

Nguyen, L. T. T., Nguyen, T. T., Nguyen, H. N., & Bui, Q. T. P. (2020). Simultaneous determination of active compounds in *Piper betle* Linn. leaf extract and effect of extracting solvents on bioactivity. Engineering Reports, 2(10), 1–8.

Noviello, S., & Huang, D. (2019). The Basics and The Advancements In Diagnosis Of Bacterial Lower Respiratory Tract Infections. Diagnostics, 9(2), 37. https://doi:10.3390/diagnostics9020037.

Okhuarobo, A., Falodun, J. E., Erharuyi, O., Imieje, V., Falodun, A., & Langer, P. (2014). Harnessing the medicinal properties of *Andrographis paniculata* for diseases and beyond: a review of its phytochemistry and pharmacology. Asian Pacific Journal of Tropical Disease, 4(3), 213–222.

Othman, L., Sleiman, A., & Abdel-Massih, R. M. (2019). Antimicrobial activity of polyphenols and alkaloids in Middle eastern plants. Frontiers in Microbiology, 10. https://doi.org/10.3389/fmicb.2019.00911.

Paderes, N. M. & Bose, M. T. (2020). Microbicidal Activity of *Tinospora rumphii boerl*, *Anamirta cocculus*, and *Andrographis paniculata* Species on Gram Positive and Gram-Negative Organisms. IAMURE International Journal of Ecology and Conservation, 30(1), 1–1.

Permadi, A. K., Pratama, E. A., Hakim, A. L. L., Widi, A. K., & Abdassah, D. (2020). The effect of carbonyl and hydroxyl compounds addition on co2 injection through hydrocarbon extraction processes. Applied Sciences, 11(1), 159. https://doi.org/10.3390/app11010159.

Prakash, B., Shukla, R., Singh, P., Kumar, A., Mishra, P. K., & Dubey, N. K. (2010). Efficacy of chemically characterized *Piper betle* L. essential oil against fungal and aflatoxin contamination of some edible commodities and its antioxidant activity. International Journal of Food Microbiology, 142(1-2), 114–119.

Prasetyoputri, A. (2021). Detection of bacterial coinfection in COVID-19 patients is a missing piece of the puzzle in the COVID-19 management in Indonesia. ACS Infectious Diseases 2021 7(2), 203-205. https://doi:10.1021/acsinfecdis.1c00006.

Pecková, R., Doležal, K., Sak, B., Květoňová, D., Kváč, M., Nurcahyo, W., & Foitová, I. (2018). Effect of *Piper betle* on *Giardia intestinalis* infection in vivo. Experimental Parasitology, 184, 39–45.

Penecilla, G. L., & Magno C. P. (2011). Antibacterial activity of extracts of twelve common medicinal plants from the Philippines. Journal of Medicinal Plants Research, 5(16), 3975–3981.

Perez, R.H. (2021). Bacteriocin production, optimization, and utilization: facilitating the utilization of bacteriocins through molecular strategies. The Microbiology Consortium of the Philippine Society for Microbiology, Inc. https://www.youtube.com/watch?v=3RFYvZWu0X4.

Rajalakshmi, V., & Cathrine, L. (2016). Phytochemical screening and antimicrobial activity of ethanolic extract of *Andrographis paniculata*. Journal of Pharmacognosy and Phytochemistry, 5(2), 175–177. https://www.phytojournal.com/archives/2016/vol5issue2/PartC/5-1-52-798.pdf

Rekha VPB, Kollipara M, Gupta Srinivasa BRSS, Bharath Y, Pulicherla KK. 2014. A Review on *Piper betle* L.: Nature’s promising medicinal reservoir. Am J Ethnomed 1 (5): 276–289.

Sahalan, A.Z., Sulaiman, N., Mohammed, N., Ambia, K.M., & Lian, H.H. (2007) Antibacterial activity of *Andrographis paniculata* and *Euphorbia hirta* methanol extracts. Jurnal Sains Kesihatan Malaysia, 5(2), 1–8.

Schmitt, D. M., Connolly, K. L., Jerse, A. E., Detrick, M. S., & Horzempa, J. (2016). Antibacterial activity of resazurin-based compounds against Neisseria gonorrhoeae in vitro and in vivo. International journal of antimicrobial agents, 48(4), 367–372. https://doi.org/10.1016/j.ijantimicag.2016.06.009.

Schmitt, D. M., O’Dee, D. M., Cowan, B. N., Birch, J. W. M., Mazzella, L. K., Nau, G. J., & Horzempa, J. (2013). The use of resazurin as a novel antimicrobial agent against *Francisella tularensis*. Frontiers in Cellular and Infection Microbiology, 3. https://doi.org/10.3389/fcimb.2013.00093.

Siahaan, R. G. (2016). Efek Kombinasi Ekstrak Etanol Daun Sirih (*Piper betle* L) dengan Amoksisilin Terhadap Pertumbuhan Bakteri *Escherichia coli* dan *Staphylococcus aureus* [Effect of betel leaf (*Piper betle* l) ethanol extract in combination with amoxicillin on the growth of *Escherichia coli* and *Staphylococcus aureus*]. TALENTA Conference Series: Tropical Medicine (TM). http://repositori.usu.ac.id/handle/123456789/12840.

Singh, J. (2008). Maceration, percolation and infusion techniques of extraction of medicinal and aromatic plants (MAPs). India: Central Institute of Medicinal and Aromatic Plants (CIMAP) Lucknow. http://www.philadelphia.edu.jo/academics/s_telfah/uploads/method%20of%20extraction.pdf.

Sitorus, P. (2018, October). Uji Efek Kombinasi Amoksisilin Dengan Ekstrak Etanol Daun Sirih (*Piper betle* L) Terhadap Pertumbuhan Bakteri *Escherichia coli* Dan *Staphylococcus aureus* [Effect test of amoxicillin combination with betel leaf (*Piper betle* L) ethanol extract on the growth of *Escherichia coli* and *Staphylococcus aureus* Bacteria]. In Talenta Conference Series: Tropical Medicine (TM), 1(1), 313–319.

Stefanović, O. (2012). Effects of plant extracts on bacterial growth and their synergistic activity with antibiotics in vitro [Thesis]. Faculty of Science: University of Kragujevac.

Stefanovic, O. D. (2018). Synergistic activity of antibiotics and bioactive plant extracts: a study against Gram-positive and Gram-negative bacteria. Bacterial Pathogenesis and Antibacterial Control, 23, 23–48. DOI: 10.5772/intechopen.72026.

Stefanović, O. D., Stanojevic, D. D., & Comic, L. R. (2012). Synergistic antibacterial activity of Salvia officinalis and Cichorium intybus extracts and antibiotics. Acta Poloniae Pharmaceutica, 69(3), 457–463.

Sule, A., Ahmed, Q. U., Latip, J., Samah, O. A., Omar, M. N., Umar, A., & Dogarai, B. B. S. (2012). Antifungal activity of *Andrographis paniculata* extracts and active principles against skin pathogenic fungal strains in vitro. Pharmaceutical Biology, 50(7), 850–856.

Syahidah, A., Saad, C., Hassan, M., Rukayadi, Y., Norazian, M., & Kamarudin, M. (2017). Phytochemical analysis, identification and quantification of antibacterial active compounds in betel leaves, *Piper betle* methanol extract. Pakistan Journal of Biological Sciences, 20(2), 70–81. https://doi.org/10.3923/pjbs.2017.70.81.

Talaro, K. P. (2008). Foundations in Microbiology (6th ed.). McGraw-Hill Education, New York, NY. pp. 370–377.

Taukoorah, U., Lall, N., & Mahomoodally, F. (2016). *Piper betle* L. (betel quid) shows bacteriostatic, additive, and synergistic antimicrobial action when combined with conventional antibiotics. South African Journal of Botany, 105, 133–140. https://doi.org/10.1016/j.sajb.2016.01.006.

Teh, C. H., Nazni, W. A., Nurulhusna, A. H., Norazah, A., & Lee, H. L. (2017). Determination of antibacterial activity and minimum inhibitory concentration of larval extract of fly via resazurin-based turbidometric assay. BMC Microbiology, 17(1). https://doi.org/10.1186/s12866-017-0936-3.

Trouger, C., Forouzanfar, M., Rao, P. C., Khalil, I., Brown, A., Swartz, S., & Reiner, R.C. (2017). Estimates of the global, regional, and national morbidity, mortality, and aetiologies of lower respiratory tract infections in 195 countries: a systematic analysis for the Global Burden of Disease Study 2015. The Lancet Infectious Diseases, 17(11), 1133–1161. doi:10.1016/s1473-3099(17)30396-1.

Valle, D. L., Andrade, J. I., Puzon, J. J. M., Cabrera, E. C., & Rivera, W. L. (2015). Antibacterial activities of ethanol extracts of Philippine medicinal plants against multidrug-resistant bacteria. Asian Pacific Journal of Tropical Biomedicine, 5(7), 532–540. https://doi:10.1016/j.apjtb.2015.04.005.

Valle, D. L., Cabrera, E. C., Puzon, J. J. M., & Rivera, W. L. (2016). Antimicrobial activities of methanol, ethanol and supercritical co2 extracts of Philippine *Piper betle* L. on clinical isolates of gram positive and gram-negative bacteria with transferable multiple drug resistance. PLOS ONE, 11(1), 1–14. https://doi.org/10.1371/journal.pone.0146349.

Xie, Y., Yang, W., Tang, F., Chen, X., & Ren, L. (2014). Antibacterial activities of flavonoids: structure-activity relationship and mechanism. Current Medicinal Chemistry, 22(1), 132–149. https://doi.org/10.2174/0929867321666140916113443

Zaidan, M. R., Noor Rain, A., Badrul, A. R., Adlin, A., Norazah, A., & Zakiah, I. (2005). In vitro screening of five local medicinal plants for antibacterial activity using disc diffusion method. Tropical Biomedicine, 22(2), 165–170.

